# Phosphorylation patterns modulate the transient secondary structure of RNA polymerase II CTD without altering its global conformation

**DOI:** 10.1101/2025.01.08.631975

**Authors:** Wei Chen (陳瑋), Tatiana N. Laremore, Neela H. Yennawar, Scott A. Showalter

## Abstract

The intrinsically disordered C-terminal domain (CTD) of RNA polymerase II coordinates transcription and co-transcriptional events through dynamic phosphorylation patterns. While it has been long hypothesized that phosphorylation induces structural changes in the CTD, a direct comparison of how different phosphorylation patterns modulate the CTD conformation has been limited. Here, we generated two distinct phosphorylation patterns in an essential *Drosophila* CTD region with the kinase Dyrk1a: one where Ser2 are primarily phosphorylated, mimicking the state near transcription termination, and a hyperphosphorylation state where most Ser2, Ser5, and Thr4 residues are phosphorylated, expanding on our work on Ser5 phosphorylation, which mimics early transcription elongation. Using ^13^C Direct-Detect NMR, we show that the CTD has a tendency to form transient beta strands and beta turns, which is altered differently by Ser2 and Ser5 phosphorylation. Small angle x-ray scattering (SAXS) revealed no significant changes in the CTD global dimensions even at high levels of phosphorylation, contradicting the common assumption of phosphorylation-induced chain expansion. Our findings support a transient beta model in which unphosphorylated CTD adopts transient beta strands at Ser2 during transcription pre-initiation. These transient structures are disrupted by Ser5 phosphorylation in early elongation, and later restored by Ser2 phosphorylation near termination for recruiting beta turn-recognizing termination factors.

## Introduction

The intrinsically disordered C-terminal domain (CTD) of RNA polymerase II regulates the transcription cycle and co-transcriptional RNA processing through its dynamic phosphorylation patterns (1–4). The CTD comprises tandem repeats of the consensus heptad sequence Y_1_S_2_P_3_T_4_S_5_P_6_S_7_, with the number of repeats varying from 26 in yeast to 52 in vertebrates (2). During transcription, CTD heptads undergo constant phosphorylation and dephosphorylation, creating distinct phosphorylation patterns – collectively known as the CTD code (5). These phosphorylation patterns specify the progress of transcription and recruit various regulators including the Mediator complex, mRNA processing enzymes, and termination factors (2, 6–9). Genome-wide analyses revealed a universal CTD code across genes (1), with phosphorylation at Ser5 and Ser2 being the most abundant (10, 11). Phospho-Ser5 (pSer5) levels peak early in transcription elongation and decline progressively, while phospho-Ser2 (pSer2) levels increase during elongation and peak near termination (1, 12). Phospho-Thr4 (pThr4), although less abundant (10, 11), is also enriched near termination (13). It has been proposed that phosphorylation patterns modulate the conformation of the CTD in distinct ways to mediate interactions with various co-regulators for co-transcriptional RNA processing (14).

Structural characterization of the CTD has been challenging due to its intrinsically disordered nature. Studies on synthetic CTD peptides with nuclear magnetic resonance (NMR) and circular dichroism (CD) spectroscopy revealed its lack of structure and a weak propensity toward beta turn (<10%) and polyproline II (<15%) (15–17). Beta turns form around Ser2 and Ser5 and are stabilized in trifluoroethanol or in circular peptides (16, 18). Because of the low population and the transient nature of secondary structures in the CTD, structural changes associated with phosphorylation are also difficult to probe. NMR and CD spectroscopy were not able to detect stable beta turns in CTD phospho-peptides (17, 19). However, when the CTD is bound to binding partners, its beta turns become stabilized and were captured in a number of crystal structures in both unphosphorylated and phosphorylated states (7, 20–27). While IDPs with alpha helical propensities and their roles in molecular recognition have been extensively studied (28–30), literature on IDPs with transient beta strands and beta turns remains limited. The CTD provides a critical example for exploring the structural and functional roles of transient secondary structures beyond alpha helices.

Phosphorylation of the CTD has been assumed to expand the global conformation of the CTD as a result of electrostatic repulsion between negatively charged phosphate groups (3, 4, 14, 16, 31). Similar expansion is expected for other IDPs that undergo multisite phosphorylation (32). This assumption is supported by studies using electrophoresis, gel filtration chromatography, sucrose gradient ultracentrifugation, and limited proteolysis (33–35), where unphosphorylated and phosphorylated CTD behave differently. However, small-angle x-ray scattering (SAXS) studies on a 12-heptad region of *Drosophila* CTD and MBP-tagged full-length *Drosophila* CTD suggest minimal changes in the radius of gyration (Rg) upon phosphorylation (34, 36).

How different phosphorylation patterns impact the CTD conformation throughout transcription remains unclear. Although several NMR and computational studies have explored the conformational properties of short CTD peptides (typically 1-3 heptads) with different phosphorylation states (19, 31, 37), structural characterization of physiologically relevant, longer phosphorylated CTDs remains limited (34, 36). In our previous work, we addressed this gap by focusing on the CTD of *Drosophila melanogaster* (34, 36). The *Drosophila* CTD consists of 42 heptad repeats, only two of which follow the exact consensus sequence Y_1_S_2_P_3_T_4_S_5_P_6_S_7_, allowing for residue-specific characterization with NMR and mass spectrometry. Previously, we characterized a 12-heptad *Drosophila* CTD region essential for viability, referred to as CTD2’, in unphosphorylated and pSer5 states, and discovered that pSer5 promotes *cis*-Pro6 formation in heptads containing Asn7 (36). This work provides insights into the structural consequences of a phosphorylation state similar to the state in early elongation, but structural effects of other important phosphorylation marks, particularly pSer2, remain largely unexplored.

Here, we investigate the conformational properties of CTD2’ with two new phosphorylation patterns created by the kinase Dyrk1a using ^13^C Direct-Detect NMR and small-angle x-ray scattering (SAXS). The phosphorylation patterns include one where Ser2 and Thr4 are primarily phosphorylated, mimicking the state near transcription termination, as well as a hyperphosphorylation state, where all Ser2, Ser5, as well as Thr4 and other non-consensus Thr residues are phosphorylated. These additional phosphorylation patterns allow for a direct comparison of structural changes induced by pSer5 vs. pSer2, providing insights for the structural consequences of changing phosphorylation patterns during transcription. We found that the global dimensions of CTD2’ remain unchanged even when 20% of the residues are phosphorylated. Phosphorylation at Ser5 leads to a decrease in the beta turn propensity while phosphorylation at Ser2 does not. Our data support a transient beta model in which transient beta turns in unphosphorylated CTD pre-initiation are disrupted by Ser5 phosphorylation in early elongation, and are restored by Ser2 phosphorylation near termination, facilitating termination factor recruitment.

## Results

### Dyrk1a preferentially phosphorylates Ser2 and Thr4 in CTD2’

In our previous study, we phosphorylated CTD2’ using the positive transcription elongation factor P-TEFb, which generated up to 12 phosphorylation marks, predominantly at Ser5 residues (36, 38). To investigate how different phosphorylation patterns affect the conformational properties of the CTD, we expanded our CTD kinase toolkit to include Dyrk1a (dual specificity tyrosine-phosphorylation-regulated kinase 1A), a kinase associated with Down syndrome that has been reported to phosphorylate Ser2 specifically *in vitro* (39–41). To generate a distinct phosphorylation pattern from that generated by P-TEFb, we performed *in vitro* phosphorylation of CTD2’ with Dyrk1a and analyzed the results using mass spectrometry (MS) and NMR.

To quantify the degree of phosphorylation, we performed MALDI-TOF MS experiments on intact CTD2’ phosphorylated by Dyrk1a for various incubation times (**Figure 1A**). Since CTD2’ contains 12 heptad repeats, 10 of which include a Ser2 residue, complete phosphorylation at every Ser2 but not at other positions would yield a mass shift of +800. However, we detected more than 10 phosphorylation events after 1 hour of incubation, indicating that Dyrk1a is not strictly specific to Ser2 in CTD2’. Incubation for 24 hours resulted in up to 19 phosphorylation events with no further phosphorylation observed after 48 hours (**Figure S1A**).

**Figure 1.**
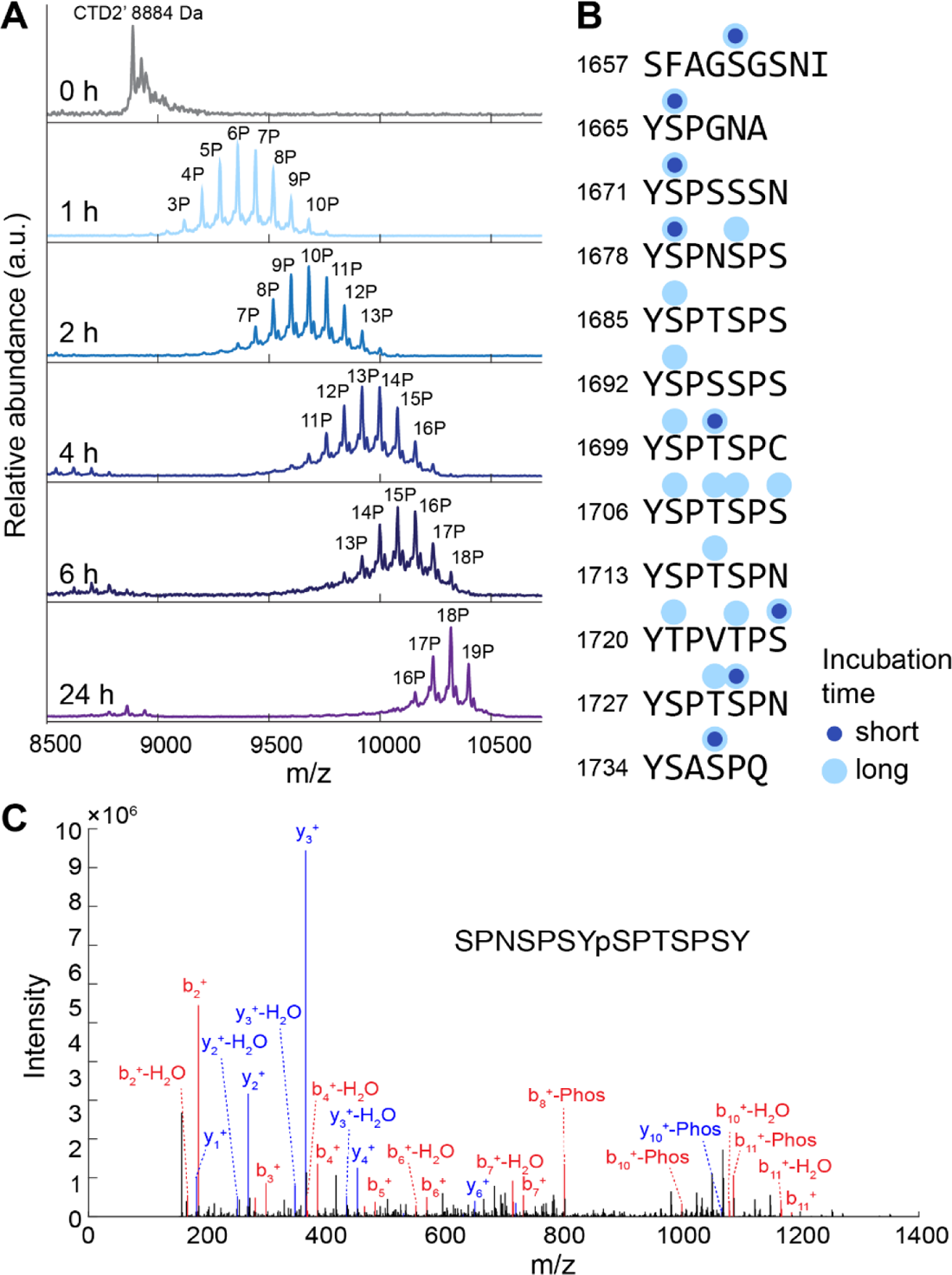
Phosphorylation of CTD2’ by Dyrk1a characterized with mass spectrometry. **(A)** MALDI-TOF mass spectra of intact CTD2’ after phosphorylation by Dyrk1a for varying incubation times. **(B)** Phosphorylation sites identified by LC-MS2 analysis of Dyrk1a phosphorylated CTD2’ with different incubation times. **(C)** Representative peptide spectrum from LC-MS2 analysis of Dyrk1a phosphorylated CTD2’. The high degree of phosphorylation suggested that Dyrk1a also phosphorylated additional Ser/Thr residues. Previous studies reporting the *in vitro* specificity of Dyrk1a for Ser2 were conducted using CTD constructs with only consensus heptads (YSPTSPS). To investigate the specificity of Dyrk1a toward CTD2’, which contains only two consensus heptads among its 12 repeats, we performed liquid chromatography-tandem mass spectrometry (nano-LC MS^2^) on chymotrypsin-digested phosphorylated CTD2’. Phosphorylation sites were identified for two phosphorylated CTD2’ samples incubated with Dyrk1a for different durations, corresponding to an average of 6 and 12 phosphorylation marks, respectively as quantified by MALDI-TOF (**Figure S1B, C**). Both data sets had 100% sequence coverage for LC-MS2 (**Figure S1B, C**), and showed that Dyrk1a preferentially phosphorylated Ser2 over Ser5 or Ser7 (Figure 1B**, C**), although phosphorylation at Ser5 and Ser7 was also observed. Interestingly, Thr residues were phosphorylated to a significant degree, which has not been previously reported for Dyrk1a(39, 40). In the first consensus heptad (residues 1685 to 1691), only Ser2 was phosphorylated, whereas in the second consensus heptad (residues 1706 to 1712), all Ser and Thr were phosphorylated.

To cross-examine the phosphorylation sites identified by MS, we conducted NMR analysis on Dyrk1a phosphorylated CTD2’ samples containing an average of 8 phosphorylation marks and a fully phosphorylated state containing 19 marks (**Figure S2A**). These samples represent two distinct states: one where, each heptad repeat contains approximately a single phosphorylated residue, and a hyperphosphorylated state where all Dyrk1a target sites are modified. The ^1^H, ^15^N-HSQC spectra of all CTD2’ samples exhibited severe signal overlap, particularly in the ^1^H dimension (Figure 2A), which is common for IDPs. This lack of spectral dispersion was especially pronounced for CTD2’ due to its highly repetitive sequence. Similar crowded ^1^H, ^15^N-HSQC spectra were observed in our previous studies of P-TEFb phosphorylated CTD2’ (36).

**Figure 2.**
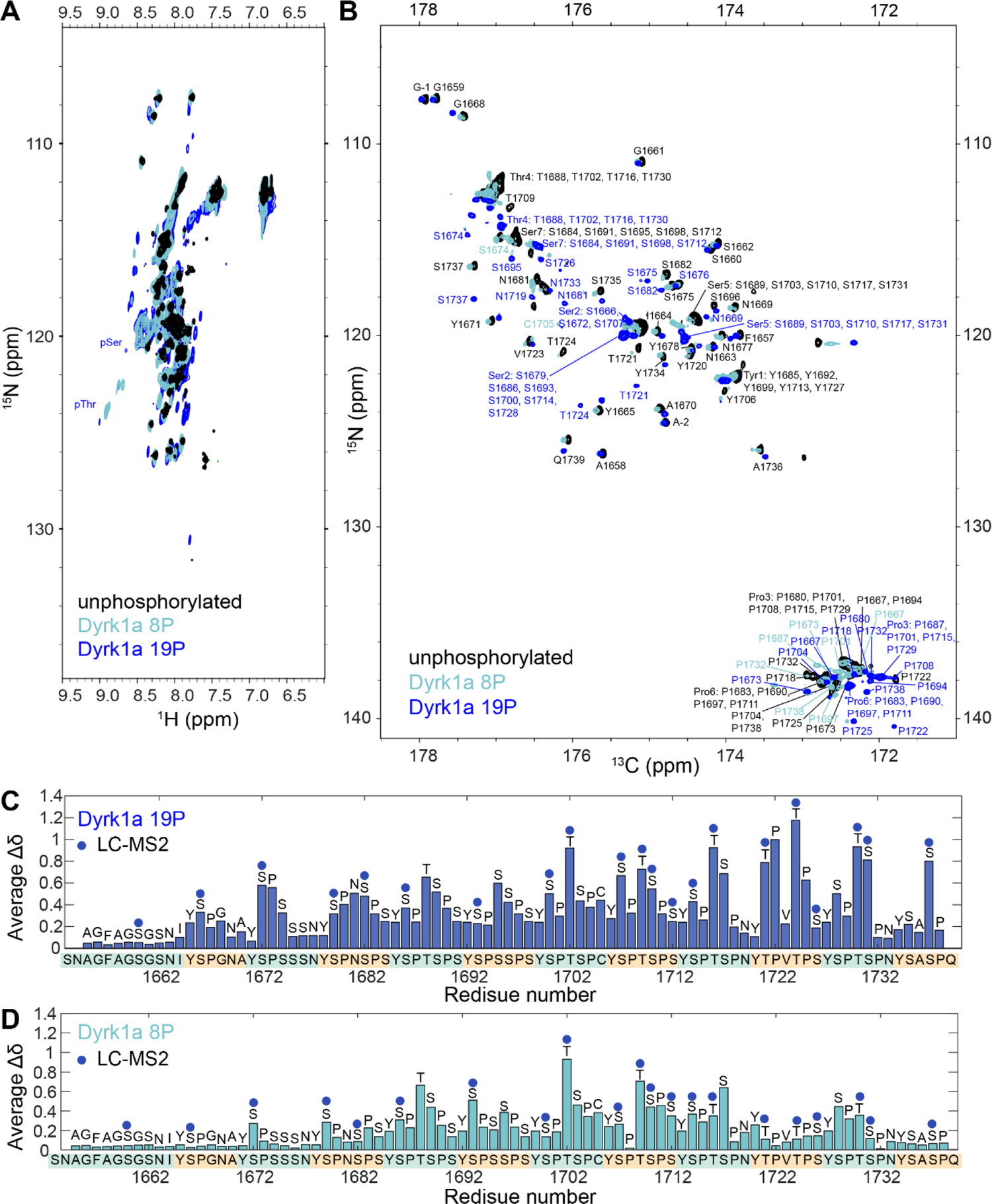
Phosphorylation of CTD2’ by Dyrk1a characterized with NMR. **(A)** 1H, 15N-HSQC spectra **(B)** CON spectra of unphosphorylated CTD2’ and Dyrk1a phosphorylated CTD2’ with 8 and 19 phosphorylation marks (8P and 19P). **(C)** Chemical shift perturbation (CSP) analysis for CTD2’ 19P reveals phosphorylation at Ser and Thr residues. **(D)** CSP analysis of CTD2’ 8P shows an enrichment of pSer2 and pThr4.

While downfield shifts in the ^1^H dimension for Ser and Thr residues upon phosphorylation were observed (Figure 2A), complete residue-level assignment for CTD2’ based on ^1^H, ^15^N-HSQC was not feasible due to signal overlap. Additionally, the abundance of Pro residues in CTD2’, which lack amide protons and are therefore invisible in ^1^H, ^15^N HSQC, further complicated conventional proton-detected NMR assignment methods. To overcome these challenges, we utilized ^13^C Direct-Detect NMR, as in our previous CTD2’ studies (36). We acquired two-dimensional CON spectra for unphosphorylated and Dyrk1a phosphorylated CTD2’ (Figure 2B). The CON spectra demonstrated remarkable spectral dispersion for all CTD2’ samples and allowed for the direct detection of Pro residues. Using a suite of two-dimensional and three-dimensional ^13^C Direct-Detect NMR experiments (42), we successfully assigned the CTD2’ spectra.

We then examined chemical shift perturbation (CSP) caused by phosphorylation. For the hyperphosphorylated CTD2’ sample containing 19 phosphorylation marks (referred to as 19P), the Ser and Thr residues showing significant CSPs were consistent with the phosphorylation sites identified by nano-LC MS^2^ (Figure 2C). Since the 19P NMR sample had more phosphorylation marks than the MS sample, additional phosphorylation at T1688 (Thr4), S1689 (Ser5), S1717 (Ser5), and S1695 was observed. This hyperphosphorylation pattern was enriched in pSer and pThr residues adjacent to Pro residues, leading to increased levels of pSer2, pSer5, and pThr. The CON spectrum of the CTD2’ sample with an average of 8 phosphorylation marks (referred to as 8P) displayed multiple sets of peaks. Some peaks overlapped with those from the unphosphorylated and hyperphosphorylated CTD2’, while others were unique (**Figure S2B**), indicating a mixture of phosphorylation states. CSP analysis of the most intense peaks in the CTD2’ 8P spectrum, which represent the major species in the mixture, showed fewer phosphorylation sites compared to the 19P sample as expected, with an enrichment of pSer2 and pThr4 (Figure 2D). Notably, this pattern enriched in pSer2 and pThr4 resembles the *in vivo* CTD phosphorylation pattern near transcription termination, in contrast to the pattern generated by P-TEFb, which mirrors the *in vivo* phosphorylation pattern near the transcription start site.

### Pro residues remain in trans states in Dyrk1a phosphorylated CTD2’

We previously showed that phosphorylation at Ser5 in CTD2’ promotes the formation is *cis*-Pro6 in the non-consensus heptad Y_1_S_2_P_3_T_4_S_5_P_6_N_7_. Since Dyrk1-phosphorylated CTD2’ contains a heptad with the sequence Y_1_S_2_P_3_N_4_S_5_P_6_S_7_, where Ser2 (S1679) is in the context of SPN, we examined whether pSer2 in the SPN motif can also induce *cis*-Pro formation in CTD2’ 8P. Additionally, we investigated the Pro *cis/trans* state in CTD2’ 19P, where Ser and Thr adjacent to Pro residues, including both Ser2 and Ser5, were phosphorylated. The *cis/trans* state of Pro residues were assessed using the C_β_ and C_γ_ chemical shifts from three-dimensional CCCON spectra (Figure 3). Unlike P-TEFb-phosphorylated CTD2’, we did not observe enrichment of *cis*-Pro3 in the SPN sequence context, suggesting that pSer-induced proline isomerization depends on sequence context beyond these three residues. In CTD2’ 19P, we also did not find enrichment for *cis*-Pro3 or *cis*-Pro6, suggesting that the propensity *cis*-Pro induced by Ser5 is counteracted when the Ser2 and/or other Thr are also phosphorylated.

**Figure 3.**
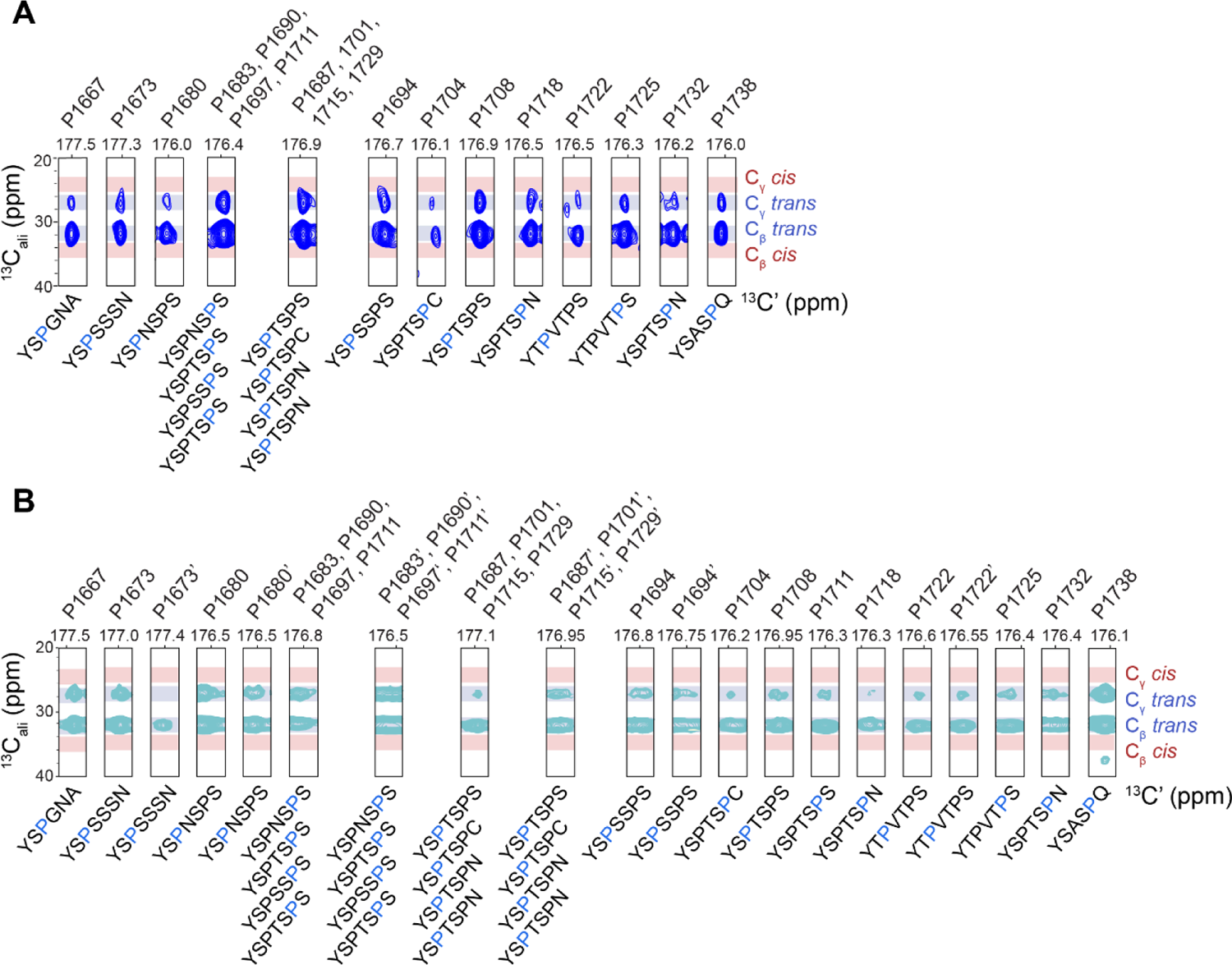
Analysis of Proline *cis-trans* conformations in Dyrk1a phosphorylated CTD2’. **(A)** C_β_ and C_γ_ chemical shifts of individual Pro residues from 3D CCCON spectra of hyperphosphorylated CTD2’ (19P). **(B)** C_β_ and C_γ_ chemical shifts of individual Pro residues from partially phosphorylated CTD2’ (8P). Blue and red shapes represent the chemical shift ranges for *trans* and *cis* Pro conformations, respectively. For partially phosphorylated CTD2’ (8P), multiple sets of peaks were observed for certain residues, with the additional set labeled with an apostrophe (‘).

### Phosphorylation patterns modulate secondary structure propensity in CTD2’

To explore how different phosphorylation patterns influence the transient secondary structure of CTD2’, we first calculated secondary structure populations using C_α_, C_β_ and backbone chemical shifts for unphosphorylated CTD2’ with the δ2D program, which has been widely used for IDPs (Figure 4A). Overall, CTD2’ adopts a coil conformation with a 23.1% extended beta strand population. Particularly, beta strand populations peak periodically at Ser2 (40%). Since δ2D does not predict beta turn populations, we used the MICS (motif identification from chemical shifts) program to estimate beta turn propensity (44). While MICS was originally trained on databases of structured proteins, it provided consistent prediction for CTD2’ with a predominantly structural disorder and a significant beta strand propensity (**Figure S3**).

**Figure 4.**
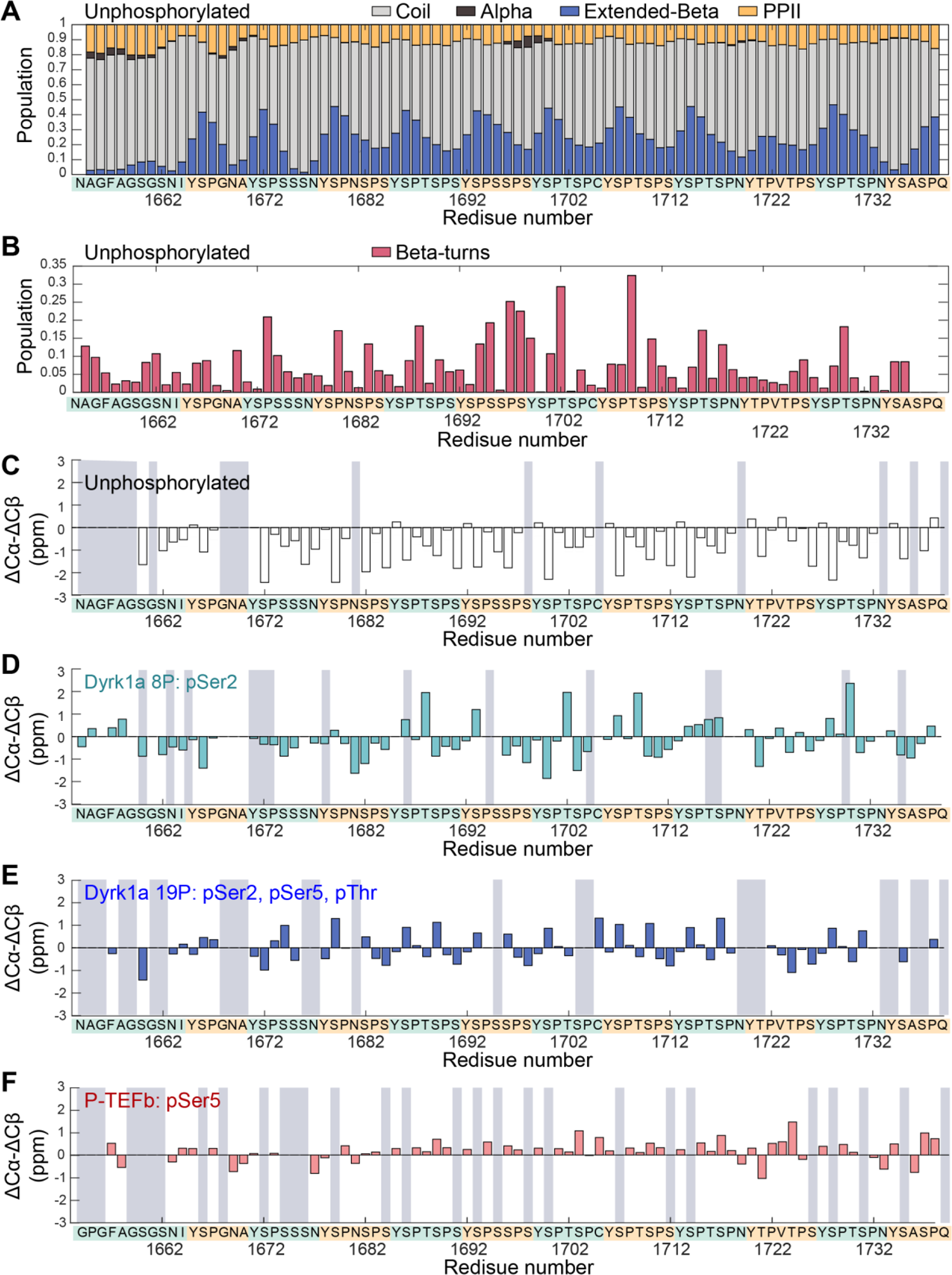
Secondary chemical propensity for CTD2’ with different phosphorylation patterns. **(A)** Populations of coil, alpha helix, extended beta strand, and polyproline II (PPII) for unphosphorylated CTD2’ calculated from backbone and sidechain chemical shifts using the δ2D program. **(B)** Population of beta turns in unphosphorylated CTD2’ calculated using the MICS program. **(C)** Secondary chemical shifts for unphosphorylated CTD2’, **(D)** Secondary chemical shifts for Dyrk1a phosphorylate CTD2’ (8P), in which Ser2 residues are primarily phosphorylated **(E)** Secondary chemical shifts for unphosphorylated for Dyrk1a phosphorylated CTD2’ (19P), in which Ser2, Ser5, and Thr residues are phosphorylated. **(F)** Secondary chemical shifts for unphosphorylated for P-TEFb phosphorylated CTD2’, in which Ser5 residues is phosphorylated. Residues without available Cα and/or Cβ chemical shift values are indicated by grey areas.

Remarkably, it identified beta turn populations at Pro3 and Thr4 across most heptads (Figure 4B**, Figure S3**), consistent with positions in NOE-based NMR studies (16, 18) and crystal structures of CTD peptides bound to Pcf11 (7). The results of δ2D and MICS align with a structural model where Tyr1 and Ser2 form transient beta strands, followed by a turn at Pro3 and Thr4. To compare the secondary structure propensity across different phosphorylation patterns, we calculated deviations of C_α_ and C_β_ chemical shifts from their expected random coil values.

Secondary chemical shifts (45), ΔC_α_-ΔC_β_, were calculated for unphosphorylated CTD2’, two Dyrk1a phosphorylated CTD2’ samples, and previously published P-TEFb phosphorylated CTD2’ (36). Positive ΔC_α_-ΔC_β_ values indicate a tendency toward alpha helix, while negative values suggest a propensity toward beta strand or extended structures. Consistent with δ2D, MICS, and our previous work (36), unphosphorylated CTD2’ shows negative ΔC_α_-ΔC_β_ values in general, indicating a disordered state with a preference for extended beta strand (Figure 4B). For phosphorylated samples, random coil chemical shift values were determined using the database published by the Kragelund group (46). For the Dyrk1a-phosphorylated CTD2’ 8P, although exact phosphorylation sites cannot be definitively assigned from the NMR spectra, a general enrichment of pSer2 was observed. We used several possible phosphorylation assignments to generate three different sets of random coil chemical shifts references, all of which produced similar results in terms of secondary chemical shifts (**Figure S4**). For Dyrk1a-phosphorylated CTD2’ 8P, ΔC_α_-ΔC_β_ values are also negative in general (Figure 4B). Compared to unphosphorylated CTD2’, the beta propensity slightly decreases for some regions, but is generally retained. For hyperphosphorylated CTD2’ 19P, where Ser2, Ser5, and Thr residues are phosphorylated, the beta propensity greatly decreases across all heptads (Figure 4C).

Interestingly, P-TEFb-phosphorylated CTD2’ (Figure 4D), which contains an average of 9 phosphorylation marks at Ser5, showed a similar degree of reduction in beta propensity as hyerphosphorylated CTD2’ 19P, which contains about twice as many phosphorylation marks. This suggests that the secondary structure propensity in CTD2’ is modulated by specific locations and patterning of phosphorylated residues rather than by the total number of phosphorylation marks. The beta propensity in unphosphorylated CTD2’ is retained when Ser2 and Thr4 become phosphorylated, but is disrupted when Ser5 residues are further phosphorylated. The beta propensity is also disrupted when phosphorylation is only at Ser5.

### CTD2’ retains its global dimensions regardless of phosphorylation states

It has been widely proposed that multi-site phosphorylation of IDPs can lead to chain expansion due to electrostatic repulsion between negatively charged phosphate groups (32). In our previous work, we showed that phosphorylation of CTD2’ by P-TEFb, which introduces an average of 9 phosphorylation marks (∼10% of the residues), did not result in changes in the radius of gyration (R_g_) and pairwise distance distribution, P(r), as determined by SAXS (36). We attributed this to Debye screening from the ions in solution. Here, we further investigate whether a higher degree of phosphorylation by Dyrk1a, which adds more closely spaced phosphate groups on Ser and Thr residues, could cause chain expansion due to expected shorter distances between charges.

We first carried out size-exclusion chromatography with multi-angle light scattering (SEC-MALS) to confirm that CTD2’ was monomeric (**Figure S5**), with a hydrodynamic radius (R_h_) of 16.12 ± 0.28 Å. We then performed SAXS on unphosphorylated and Dyrk1a-phosphorylated CTD2’ containing an average of 17 phosphorylation marks (∼20% of the residues) (**Figure S6, Table S1**). The R_g_ values calculated using the Guinier approximation are 27.99 ± 0.28 Å for unphosphorylated and 28.24 ± 0.25 Å for Dyrk1a phosphorylated CTD2’, consistent with our previously reported values (28.0 ± 0.7 for unphosphorylated and 28.3 ± 0.3 for P-TEFb-phosphorylated CTD2’). The R_g_/R_h_ ratio is typically 3/5 (∼0.775) for globular proteins, 1.24 for ideal Gaussian chains, and 1.56 for excluded volume chains (47). For CTD2’, the R_g_/R_h_ ratio is 1.736, indicating a more extended conformation compared to excluded volume chains.

According to the polymer scaling law parametrized for IDPs (48), *R_g_ = R_0_N^ν^*, in which *R_0_* and *ν* are 2.54 and 0.522 respectively, an IDP with the same number of residues as CTD2’ would have an R_g_ of 26.14 Å. This suggests that CTD2’ is slightly more extended than typical IDPs. In addition to the similarity in R_g_ values between unphosphorylated and highly phosphorylated CTD2’, there were no significant differences in the P(r) plots between the two states (Figure 5A).

**Figure 5.**
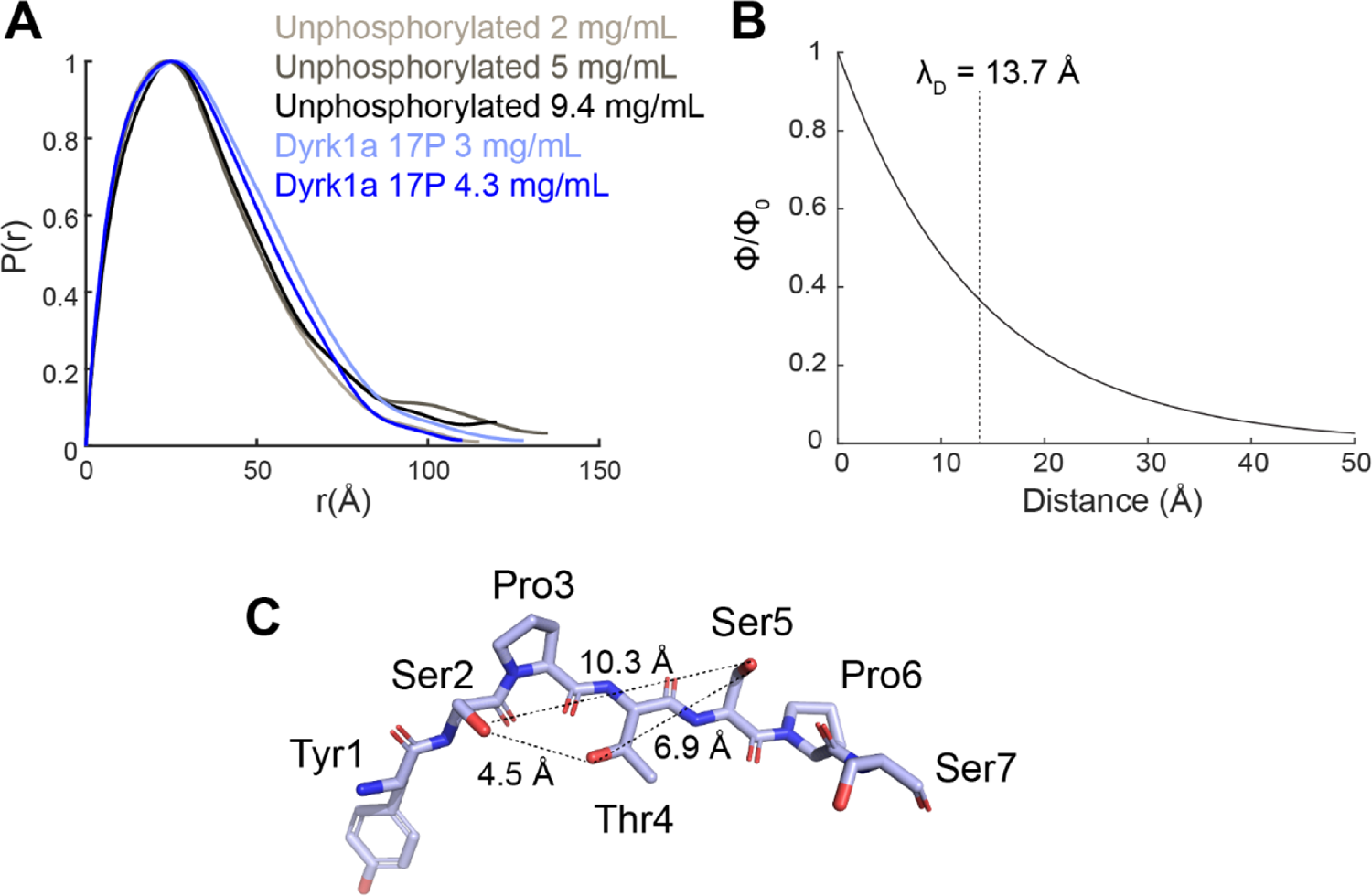
SAXS analysis of global dimensions and Debye screening in CTD2’. **(A)** Representative pair-wise distance distributions, P(r), for unphosphorylated CTD2’ (black) and Dyrk1a phosphorylated CTD2’ 17P (blue). **(B)** Debye screening of the electric potential for the SAXS buffer condition with a Debye length (λ_D_) of 13.7 Å. **(C)** Distances between Ser2, Thr4, and Ser5 residues in an extended conformation.

To evaluate whether Debye screening plays a major role in maintaining the chain dimension of highly phosphorylated CTD2’, we calculated the screening effect of electric potential using the buffer conditions from our SAXS experiments (Figure 5B). The characteristic Debye screening length (λ_D_), defined as 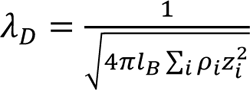, where *l_B,_* is the Bjerrum length (7 Å), was calculated as 13.7 Å for our buffer containing 50 mM KCl (49) (8.4 Å for 25 mM sodium phosphate, 50 mM KCl). The Debye screening effect decays exponentially as the distance increases (Figure 5B). To estimate the distances between phosphorylated residues in CTD2’, we measured the distances between sidechain oxygen atoms in the residue pairs, Ser2-Ser5, Ser2-Thr4, and Thr4-Ser5, in a completely extended conformation (Figure 5C). The measured distances were 10.3 Å, 4.5 Å, and 6.9 Å, respectively. Given the compact nature of CTD2’ suggested by SAXS, the actual distances between phosphorylated residues may be shorter that those measured for an extended conformation. Based on the above estimation, electrostatic interactions between phosphorylated residues in Dyrk1a phosphorylated CTD2’ do not seem to be completely screened out. This suggests that besides Debye screening, local conformational changes, as such changes in secondary structure propensity indicated by NMR, also contribute to maintaining the overall size and shape of CTD2’ upon phosphorylation.

## Discussion

The CTD of RNA polymerase II undergoes dynamic phosphorylation throughout the transcription cycle and recruits co-regulators that recognize specific phosphorylation. It has long been suggested that phosphorylation causes the CTD to adopt a more extended conformation due to charge repulsion between phosphate groups (3, 4, 14). This hypothesis has been supported by earlier observations of differences between unphosphorylated and phosphorylated CTDs in gel filtration chromatography and sucrose gradient ultracentrifugation (33), as well as a recent computational study on a 44-residue human CTD region (31). However, our previous SAXS studies on CTD2’ and the full-length *Drosophila* CTD phosphorylated at Ser5 by P-TEFb showed only modest or no changes in R_g_ (34, 36). Here, we further demonstrated that CTD2’ retains its global conformational compactness even when 20% of the total residues are phosphorylated This suggests that the CTD retains its global dimensions during the dynamic phosphorylation and dephosphorylation process in transcription, where typically ∼5% of residues are phosphorylated at a given time (10, 11).

Comparison of R_g_ values for CTD2’ (28 Å) and average IDPs with the same number of residues (24-26 Å) (51, 52) suggests that although the CTD has been described as “compact” compared to a fully extended chain, which is not thermodynamically favored, it is slightly more extended than average IDPs. Debye screening plays a role at physiological conditions (λ_D_∼7 Å) in screening out most charge interactions. However, our 20% phosphorylated CTD2’ with closely spaced phosphate groups that can have separations smaller than the Debye length suggests that the conservation of global dimension may be an inherent property coded in the amino acid sequence of the CTD. Possible conformational changes that counteract the effect of charge repulsion include compaction at the scale of 1-2 heptads as a result of beta turn formation upon phosphorylation and coordination of phosphate groups with counterions in the buffer, both of which have been observed in computational studies (31, 37). The conformational rigidity of Pro residues, as well as their *cis-trans* isomerization coupled with phosphorylation, may also contribute to offsetting the structural impact of charge repulsion. From the perspective of function, the amino acid sequence of the CTD may be evolved for it to retain the global dimension in various phosphorylated states, so that the accessibility remains the same for unphosphorylated or differently phosphorylated CTD to bind to co-regulators.

Early NMR studies on short synthetic CTD peptides showed that the CTD is highly flexible with a tendency to form beta-turns (15, 16). NMR secondary structure propensity analyses on long CTD constructs from our group and others are consistent with this view, showing that the CTD adopts a conformational ensemble with a beta-turn propensity around Ser2 (36, 53). It has been proposed for that the CTD can adopt different conformations depending on specific phosphorylation patterns (14). Specifically, a beta-spiral model has been proposed (7, 14), in which the unphosphorylated CTD adopts a compact conformation where each heptad forms a beta turn, and these turns are stacked together in a spiral-like arrangement, which is opened up by Ser5 phosphorylation but not Ser2 phosphorylation (14). Our SAXS data suggest that the CTD retains its global compactness even when highly phosphorylated. Moreover, the CTD remains disordered across all phosphorylation states and does not adopt the stable secondary structures required for building a beta spiral. Circular dichroism studies on the CTD were unable to detect population changes in beta-turns associated with phosphorylation (17, 19). With NMR chemical shifts, we are able to detect a significant degree of beta structural propensity in unphosphorylated CTD2’, which is disrupted by pSer5 but not by pSer2. Interestingly, when both Ser5 and Ser2 are phosphorylated, the beta structural propensity is also lost. Our previous work showed that pSer5 promotes *cis*-Pro6 formation in non-consensus heptads containing Asn7 (36). We note that this effect would not be unique to *Drosophila* because a similar cluster of heptads containing Asn7 is found in mammalian CTDs, including mouse and human. A computational study on single CTD heptads showed that *cis*-Pro6 inhibits beta turn formation (37), suggesting *cis*-Pro6 also plays a role in the lack of beta structure propensity for CTD2’ with pSer5.

Our NMR and SAXS data support a transient beta model in which the CTD retains its overall compactness (Figure 6), which is more extended than typical IDPs, potentially to allow accessibility to co-regulators throughout the transcription cycle, including interactions with the Mediator in the unphosphorylated state for transcription preinitiation, as well as with various mRNA processing factors in differently phosphorylated states during elongation and near termination. While the global dimensions are conserved, the changing phosphorylation patterns modulate the transient local structures of the CTD. Phosphorylation at Ser5, which is prevalent near the transcription start site, decreases the beta propensity and promotes *cis*-Pro6 formation for Asn7-containing heptads. As transcription proceeds and the level of pSer5 decreases while pSer2 increases during elongation, the beta propensity is restored and the *cis*-Pro level decreases, facilitating the recognition of beta turns and *trans*-Pro by the termination factor Pcf11 (7, 19). Notably, Pcf11 does not directly interact with the phosphate group on Ser2 but rather binds to the beta turn (7, 19). Our NMR data of the hyperphosphorylated CTD reveals that in the presence of pSer5, the beta conformational propensity remains low, despite Ser2 phosphorylation, suggesting that pSer5, in addition to pTyr1 (54), may contribute in preventing premature termination by Pcf11 binding. The beta conformation becomes sufficiently populated to recruit Pcf11 near the polyadenylation site, where Ser5 residues are mostly dephosphorylated.

**Figure 6.**
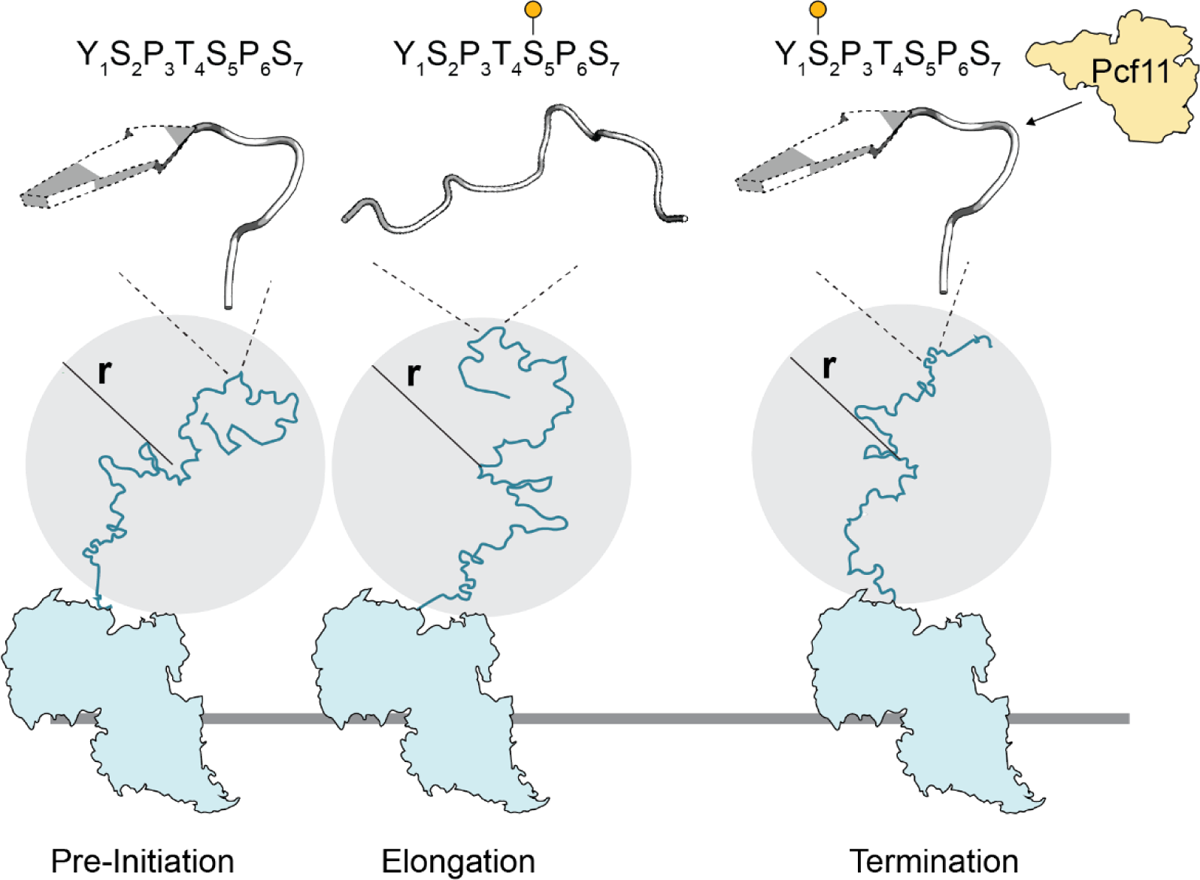
A transient beta model for CTD conformational modulation by phosphorylation. In the transcription pre-initiation state, the unphosphorylated CTD adopts a flexible conformational ensemble with a 20-40% propensity to form transient beta strands and turns around Ser2. As transcription progresses, the CTD becomes phosphorylated primarily at Ser5 near the transcription start site, which disrupts the beta turns. During elongation, the decrease of pSer5 level combined with increase of pSer2 level restores the beta propensity, which is crucial for binding to the termination factor Pcf11. Despite the changing phosphorylation patterns, the global dimensions of the CTD, as measures by r, remain conserved throughout transcription. This conservation of global size ensures accessibility to binding partners at different transcription stages, while changes in transient local structures modulate specific interactions with co-regulators.

## Conclusion

Our results show that, contrary to assumptions about phosphorylation-induced chain expansion, the CTD retains its overall compactness even in highly phosphorylated states. While the global dimensions are conversed, transient local structures are modulated by phosphorylation patterns. Phosphorylation at Ser5 inhibits beta strand formation while phosphorylation at Ser2 preserves beta propensity. This suggests that dynamic modulation of local CTD structure, rather than large-scale conformational changes, may be the key to how the CTD integrates phosphorylation signals to regulate interactions with binding partners throughout the transcription cycle.

## Materials and Methods

### Protein expression and purification

*CTD2’.* An *E. coli* codon optimized DNA sequence encoding residues 1657-1739 of the *Drosophila melanogaster* Rpb1 C-terminal domain (CTD2’) was cloned into the pET His6 MBP TEV LIC cloning vector (1M, Addgene plasmid #29656) using ligation independent cloning. This vector encodes a 6xHis-tagged maltose binding protein (MBP) and a TEV cleavage sites at the N-terminus of the CTD2’. After TEV cleavage, a cloning artifact (amino acids SNAG) remains at the N-terminus of the CTD2’, which does not affect the biophysical properties of CTD2’, as they match those of a previous construct with a GPG cloning artifact(36). BL21 (DE3) cells containing the CTD2’ plasmid were grown in LB media with 50 ug/mL kanamycin at 37 °C and 200 rpm. At an OD_600_ of 0.7-0.8, expression of was induced with 0.5 mM IPTG at 37 °C and 200 rpm for 3 hours. Cells were pelleted by centrifugation at 5000x g and stored at −80 °C. Cell pellets of CTD2’ were resuspended in the lysis buffer (50 mM Tris pH 7.5, 500 mM NaCl, 20 mM imidazole, 2.5 mM β-mercaptoenthanol) supplemented with 1x EDTA-free protease inhibitor cocktail (Millipore), 1 mM PMSF, and 10 units of RNAse-free DNAse (NEB). Following sonication on ice, the cell lysate was centrifuged at 14000x g for 45 minutes. The supernatant was passed through Ni-NTA resins (Thermo Fisher Scientific) equilibrated with the lysis buffer and washed with 10 column volumes of the lysis buffer. Proteins were eluted with the elution buffer (50 mM Tris pH 7.5, 500 mM NaCl, 200 mM imidazole, 2.5 mM β-mercaptoenthanol). The eluate was incubated with TEV protease (1 mg per 1 L growth) under dialysis against 2 L of 50 mM Tris pH 7.5, 100 mM NaCl, 2.5 mM β-mercaptoenthanol at 4 °C for 16 hours. Dialysates were passed through a second Ni-NTA column, and the flow-through containing CTD2’ was collected and concentrated using a Amicon Ultra Centrifugal Filter with a 3 kDa molecular weight cutoff (MilliporeSigma). The concentration of CTD2’ was determined by A_280_ using the extinction coefficient ε^CTD2’^ = 16390 M^-1^cm^-1^.

*Dyrk1a.* BL21 (DE3) cells were co-transformed with plamids containing the kinase domain of human Dyrk1a (Addgene Plasmid #79690) and the lambda phosphatase (Addgene Plasmid #79748)(55). Cells were grown in LB media with 100 ug/mL ampicillin and 50 ug/mL spectinomycin at 37 °C and 200 rpm until OD_600_ reaches 0.6-0.8. Expression was induced with 0.2 mM IPTG at 18 °C and 200 rpm for 16 hours. Cells were pelleted by centrifugation at 5000x g and stored at −80 °C. Cell pellets of CTD2’ were resuspended in the lysis buffer (50 mM Tris pH 7.5, 500 mM NaCl, 20 mM imidazole, 2.5 mM β-mercaptoenthanol) supplemented with 1x EDTA-free protease inhibitor cocktail (Millipore) and 1 mM PMSF. Following sonication on ice, the cell lysate was centrifuged at 14000x g for 45 minutes. The supernatant was passed through Ni-NTA resins (Thermo Fisher Scientific) equilibrated with the lysis buffer and washed with 10 column volumes of the lysis buffer. Proteins were eluted with the elution buffer (50 mM Tris pH 7.5, 500 mM NaCl, 200 mM imidazole, 2.5 mM β-mercaptoenthanol). The eluate was incubated with TEV protease (1 mg per 1 L growth) to remove the 10xHis tag under dialysis against 2 L of 50 mM Tris pH 7.5, 100 mM NaCl, 1 mM DTT at 4 °C for 16 hours. The concentration of CTD2’ was determined by A_280_ using the extinction coefficient ε^Dyrk1a^_280_ = 50310 M^-1^cm^-1^.

### Kinase reactions

Phosphorylation time course reactions using Dyrk1a were performed with 2 mL of 46 μM CTD2’, 192 μL of 49 μM Dyrk1a, 10 mM ATP, 20 mM MgCl2, and 1 mM DTT at room temperature (21°C). For MALDI-TOF analysis, 200 μL of the reaction mixture was taken at time points of 1, 2, 4, 6, and 24 hours. Each sample was heated at 90°C for 5 minutes to deactivate Dyrk1a.

### Matrix-assisted laser desorption/ionization-time of flight mass spectrometry (MALDI-TOF MS)

Samples of intact unphosphorylated and phosphorylated CTD2’ were desalted using Pierce C-18 Spin Columns (Thermo Fisher Scientific) and desiccated in a SpeedVac Vacuum Concentrator. The mass spectra were acquired on a Bruker Ultraflextreme MALDI TOF-TOF instrument using a factory-configured instrument method for the linear positive-ion detection over the 5,000 – 20,000 *m/z* range. The sample-matrix mixture was prepared by combining 1 volume of 20 mg/mL super-DHB (Sigma p/n 50862-1G-F) in 50% ACN, 0.1% trifluoroacetic acid, 0.1% phosphoric acid with 1 volume of sample, 5-10 mg/mL in water, and 1 μL of this mixture was applied to a polished steel target. Laser power attenuation and pulsed ion extraction time were optimized to achieve the best signal. Instrument was calibrated using a Bruker Protein Mix I calibration standard containing a mixture of 4 proteins: bovine insulin, bovine ubiquitin, equine heart cytochrome C, and equine apomyoglobin. Mass spectra were smoothed (SavitzkyGolay, width 5 m/z, 1 cycle) and baseline subtracted (TopHat). Mass lists were generated using a centroid peak detection algorithm.

### Nano-liquid chromatography tandem mass spectrometry (Nano-LC MS^2^)

For alkylation and chymotrypsin digestion, 100-200 ug of phosphorylated CTD2’ in 100 mM Tris-HCl pH 8, 2 mM calcium chloride, 10 mM DTT was incubated at 60 °C for 45 minutes to reduce cysteine residues. After cooling the sample to room temperature, iodoactamide (IAM) was added to a final concentration of 20 mM for cysteine alkylation. The reaction was incubated at room temperature for 30 minutes protected from light. Same amount of DTT as in the reduction step was added to quench the reaction. 1 mg/mL of Chymotrypsin Endoproteinase, TLCK treated, MS Grade (Thermo Scientific) was added to a final 1:20 enzyme-to-protein sample ratio. The reaction was incubated at 37 °C for 16-24 hours (overnight) and stored at −80 °C until mass spectrometry analysis.

The chymotryptic peptides were separated on a Thermo Easy-nLC 1200 using 0.1% formic acid in water as mobile phase A and 80% aqueous acetonitrile, 0.1% formic acid as mobile phase B at a flow rate of 300 nL/min. The peptides, 1-2 μg were loaded onto an Acclaim PepMap100 trapping column (75 μm × 2 cm, C18, 3 μm, 100 Å, Thermo) using 5% B and separated on an Acclaim PepMap RSLC column (50 μm × 15 cm, C18, 2 μm, 100 Å, Thermo) with a 30-min 5% - 25% B, followed by 10 min 25% - 45% B. The column was then flushed with 90% B for 6 min. A Thermo Eclipse mass spectrometer was operated in a cycle time dependent mode (2 s) with the following data-dependent parameters: full FT MS scan at R 120,000 followed by R 15,000 MS^2^ scans with HCD activation (30% normalized collision energy). Only the precursors with charge states 2-6 were selected for MS^2^; precursor isolation was in the quadrupole with a 1.6 *m/z* isolation window. The dynamic exclusion duration was set to 10 s.

The mass spectra were processed using the Proteome Discoverer 2.5 (Thermo) and searched with Sequest HT and MS Amanda 2.0 search engines and an IMP-ptmRS module for the phosphorylation analysis. The database contained the CTD2 and the Dyrk1a kinase sequences and 299 sequences of common contaminant proteins. The search parameters included precursor tolerance 10 ppm, fragment tolerance 0.02 Da, dynamic modifications included Phosphorylation (+79.966 Da, S, T, Y) and Oxidation (+15.995 Da, M), static modification was Carbamidomethyl (+ 57.021 Da, C). Only peptides with a High Sequest HT confidence score were used for phosphorylation site assignment.

### NMR data collection and analysis

CTD2’ for NMR experiments was expressed in M9 minimal media with ^13^C and ^15^N enrichment achieved through the incorporation of ^15^N-ammonium chloride and ^13^C D-glucose (Cambridge Isotope Laboratories). The proteins were purified as described above. Phosphorylated CTD2’ was prepared by kinase reactions with 700 μL of 434 μM CTD2’ with a 12:1 molar ratio of CTD2’ to Dyrk1a, with 10 mM ATP, 20 mM MgCl_2_, 1 mM DTT. The reaction mixture was dialyzed against 50 mM Tris pH 7.5, 100 mM NaCl, 10 mM ATP, 20 mM MgCl_2_, 1 mM DTT. Reactions were incubated for 1 hour and 48 hours at room temperature (21 °C) to produce CTD2’ with 8 and 19 phosphorylation marks, respectively. Following incubation, samples were heated at 65°C for 5 minutes and centrifuged to remove the Dyrk1a containing pellets. The heating step did not affect the NMR spectra of CTD2’.

NMR data were collected on Bruker Avance III 600 MHz or Bruker Avance NEO 600 MHz spectrometers equipped with TCI triple-resonance cyroprobes. All data collection and initial processing were carried out in TopSpin (Bruker). ^13^C chemical shifts were referenced to sodium trimethylsilylpropanesulfonate (DSS). All NMR experiments were carried out in 80 mM imidazole pH 6.5, 50 mM KCl, 2.5 mM β-mercaptoenthanol, 10% D_2_O.

Chemical shift assignments of CTD2’ were carried out using ^13^C-Direct Detect methods. Assignments were generated based on CON experiments (8 scans, 1024 (^13^C) × 256 (15N) complex data points, with sweep widths of 18 ppm and 42 ppm, respectively) collected from ∼1 mM ^13^C, ^15^N, CTD2’ samples. Three-dimensional experiments for assignments included the following: (HACA)N(CA)CON (8 scans, 1024 (^13^C) × 64 (^15^N) × 128 (^15^N), with sweep widths of 12 ppm, 42 ppm, and 42 ppm, respectively); (HACA)N(CA)NCO (8 scans, 1024 (^13^C) × 64 (^15^N) × 128 (^15^N), with sweep widths of 12 ppm, 42 ppm, and 42 ppm, respectively); CCCON (8 scans, 1024 (^13^C) × 64 (^15^N) × 128 (^13^C), with sweep widths of 12 ppm, 42 ppm, and 80 ppm, respectively). Peak picking and assignments were carried out manually using NMRFAM-Sparky. Average chemical shift perturbations were calculated as follows:

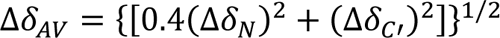

Random coil values of phosphorylated CTD2’ at 25 °C and pH 6.5 were obtained from the Kragelund database (https://www1.bio.ku.dk/english/research/bms/sbinlab/randomchemicalshifts2/)

### Size-exclusion chromatography with multi-angle light scattering (SEC-MALS)

After purification as described above, CTD2’ was further purified using size exclusion chromatography with P-10 resins in 80 mM imidazole pH 6.5, 50 mM KCl, 1mM DTT. The sample was then concentrated to 1.6 mM. 20 μL of CTD2’ was loaded onto a Superdex S-100 Increase column (Cytiva) equilibrated with 25 mM sodium phosphate pH 7.5, 50 mM NaCl, 4 mM DTT.

SEC–MALS experiment was conducted using an Agilent 1260 Infinity II HPLC system equipped with an autosampler all set to room temperature. The column used was the Superdex 200.

Wyatt Technology DAWN MALS, Wyatt Optilab refractive index detector and Agilent UV detector were used for analyzing the molar mass of peaks that eluted from the column. The SEC–MALS system was equilibrated for 5 h with buffer consisting of 25 mM sodium phosphate pH 7.5, 50 mM NaCl, 4 mM DTT. UV was set to 280nm and temperature was set to 4C. A volume of 20µl of sample was injected at a flow rate of 0.5 ml min^−1^ with a chromatogram run time of 48 min. The sample concentration was 10mg/mL. Dynamic light scattering data was collected using an inline Wyatt Dynapro Nanostar equipment.

Normalization and alignment of the MALS and refractive index detectors were carried out on standard BSA. The concentration source used was refractive index. Data were analyzed using the ASTRA software Version 8.2.2 (Wyatt). The chromatogram showed a single peak in the light scattering and the refractive index that was monodisperse. Molar mass (Mw) showed a mass consistent with a monomer = 8.8 kDa. Hydrodynamic radius was estimated to be 16.12 (±1.761%) Å.

### Small-angle x-ray scattering (SAXS)

SAXS experiments were performed using a Rigaku MM007 rotating anode X-ray source paired with the BioSAXS2000nano Kratky camera system. This setup includes OptiSAXS confocal max-flux optics, designed specifically for SAXS, and a highly sensitive HyPix-3000 Hybrid Photon Counting detector. The sample-to-detector distance was calibrated to 495.5 mm using silver behenate powder (The Gem Dugout, State College, PA). The momentum transfer scattering vector (q-space) range, defined as 4πsin(θ)/λ, extended from qmin = 0.008 Å⁻¹ to qmax = 0.6 Å⁻¹ (where q = 4πsin(θ)/λ and 2θ is the scattering angle). The X-ray beam energy was set to 1.2 keV, with a Kratky block attenuation of 23% and a beam diameter of approximately 100 μm. For sample handling, protein samples were loaded into a quartz capillary flow cell using the Rigaku autosampler, with the sample stage maintained at room temperature. The entire X-ray flight path, including the beam stop, was held under vacuum conditions of < 1×10⁻³ torr to reduce air scattering. Automated data collection, including detailed cleaning cycles between samples, was managed by the Rigaku SAXSLAB software.

After purification as described above, samples of CTD2’ for SAXS was further purified using size exclusion chromatography with P-10 resins in 80 mM imidazole pH 6.5, 50 mM KCl, 1mM DTT. Phosphorylation reactions were set up with a 60:1 molar ratio of CTD2’ to Dyrk1a, in the presence of 10 mM ATP, 20 mM MgCl_2_, 1 mM DTT at 37 °C for 40 min, followed by incubation at room temperature (21 °C) for 48 hours. Phosphorylation of CTD2’ was confirmed by MALDI-TOF mass spectrometry. Dyr1ka was removed using Q Sepharose HP chromatography in 25 mM Tris pH 7.5 and 1 mM DTT with a KCl gradient from 50 mM to 1 M. The eluted phosphorylated CTD2’ was then buffer exchanged into 25 mM Tris pH 7.5, 50 mM KCl, 1 mM DTT. For SAXS, two sample concentrations were used: 5 mg/mL and 9.4 mg/mL for unphosphorylated CTD2’, and 3 mg/mL and 4.3 mg/mL for Dyrk1a phosphorylated CTD2’. We also performed SAXS on the SEC-MALS eluted fraction of unphosphorylated CTD2’, which was at a concentration of 2 mg/mL.

Data processing was completed using the Rigaku SAXSLAB software, with each dataset averaged from six ten-minute images across three replicates of both protein and buffer samples to confirm the absence of X-ray radiation damage. Consistency in SAXS data overlays verified no radiation decay or sample loss over the 60-minute collection period. Reference buffer subtraction was subsequently performed to isolate the protein’s raw SAXS scattering curve. The forward scattering I(0) and radius of gyration (Rg) were calculated using the Guinier approximation, which assumes the intensity follows I(q) = I(0)exp[−1/3(qRg)²] at very small angles (q < 1.3/Rg). Data analysis included evaluating parameters such as radius of gyration, D_max_, Guinier fits, Kratky plots, and the pair distance distribution function with ATSAS software. The pairwise distance distribution function P(r) was generated using GNOM following protocols for IDPs (56, 57), where D_max_ was increased until P(r) smoothly approaches zero at larger r.

## Acknowledgement

We thank Julia Fecko at the Penn State X-ray Crystallography Core Facility for conducting and analyzing the SEC-MALS experiments, and Olivia Fraser for assistance in setting up the NMR experiments. This work was supported by funding from the U.S. National Science Foundation (MCB-1932730 to S.A.S.). W.C. acknowledges support from the Eberly Postdoctoral Research Fellowship. Research reported in here was also supported by SIG S10 of the National Institutes of Health under award number # S10-OD028589 for the small angle X-ray scattering and S10 OD030490 for the Wyatt SEC-MALS-DLS system to N.H.Y.

## Data Availability

NMR assignments for unphosphorylated and Dry1ka-phosphorylated (8P and 19P) CTD2’ have been deposited in the BMRB database.

