## Supplementary Information for "Phosphorylation patterns modulate the transient secondary structure of RNA polymerase II CTD without altering its global conformation"

Figures S1-S6

Table S1

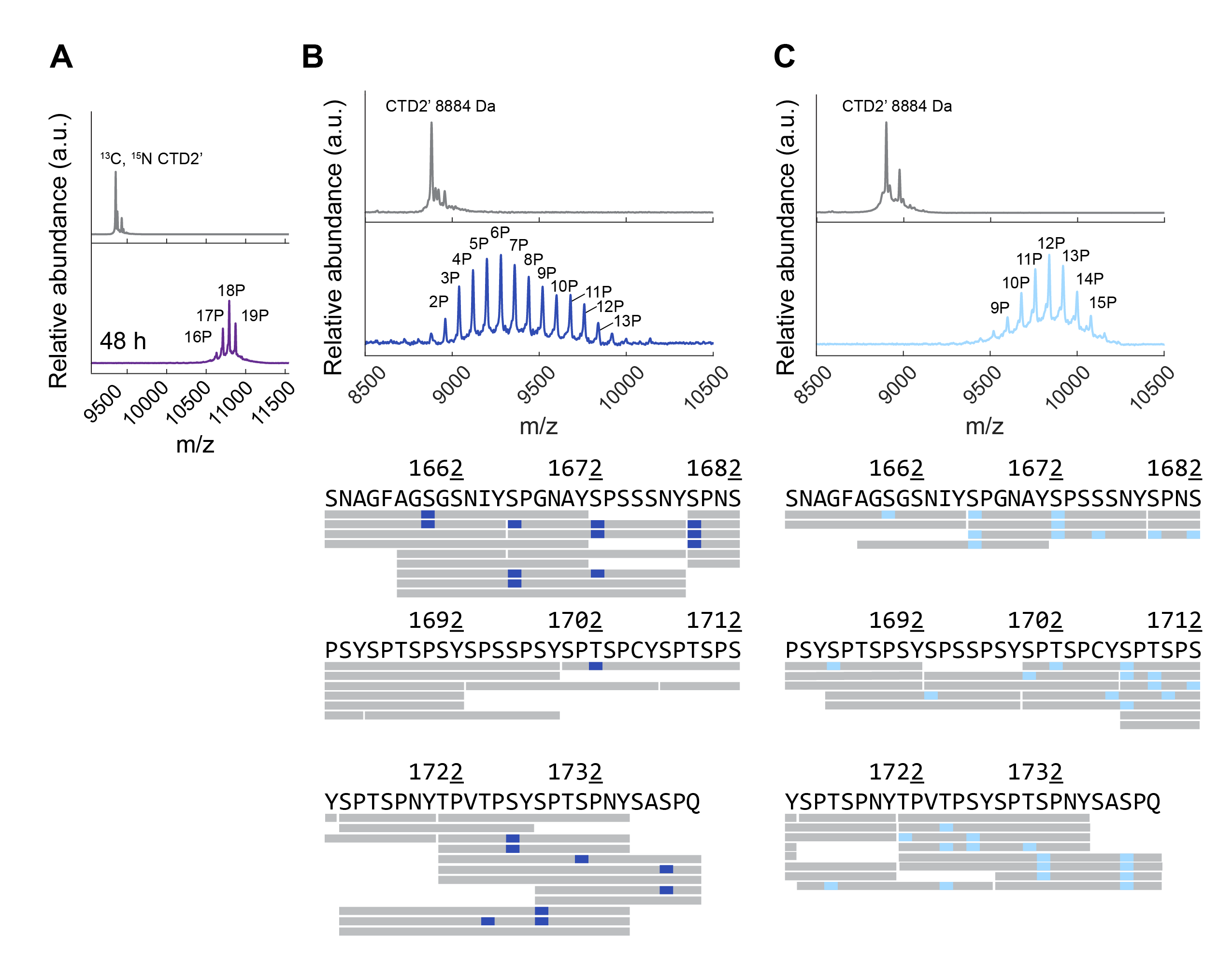

**Figure S1.** **Phosphorylation of CTD2’ by Dyrk1a characterized with mass spectrometry.** **(A)** MALTI-TOF mass spectra of unphosphorylated and Dyrk1a phosphorylated CTD2’ after 48 hours of incubation, showing a maximum of 19 phosphorylation events. No additional phosphorylation marks are observed compared to the 24-hour sample. **(B, C)** MALDI-TOF spectra of Dyrk1a phosphorylated CTD2’ with an average number of 6 and 12 phosphorylation marks, and the corresponding sequence coverage of chymotrypsin digested peptides showing phosphorylation sites identified by LC-MS2.

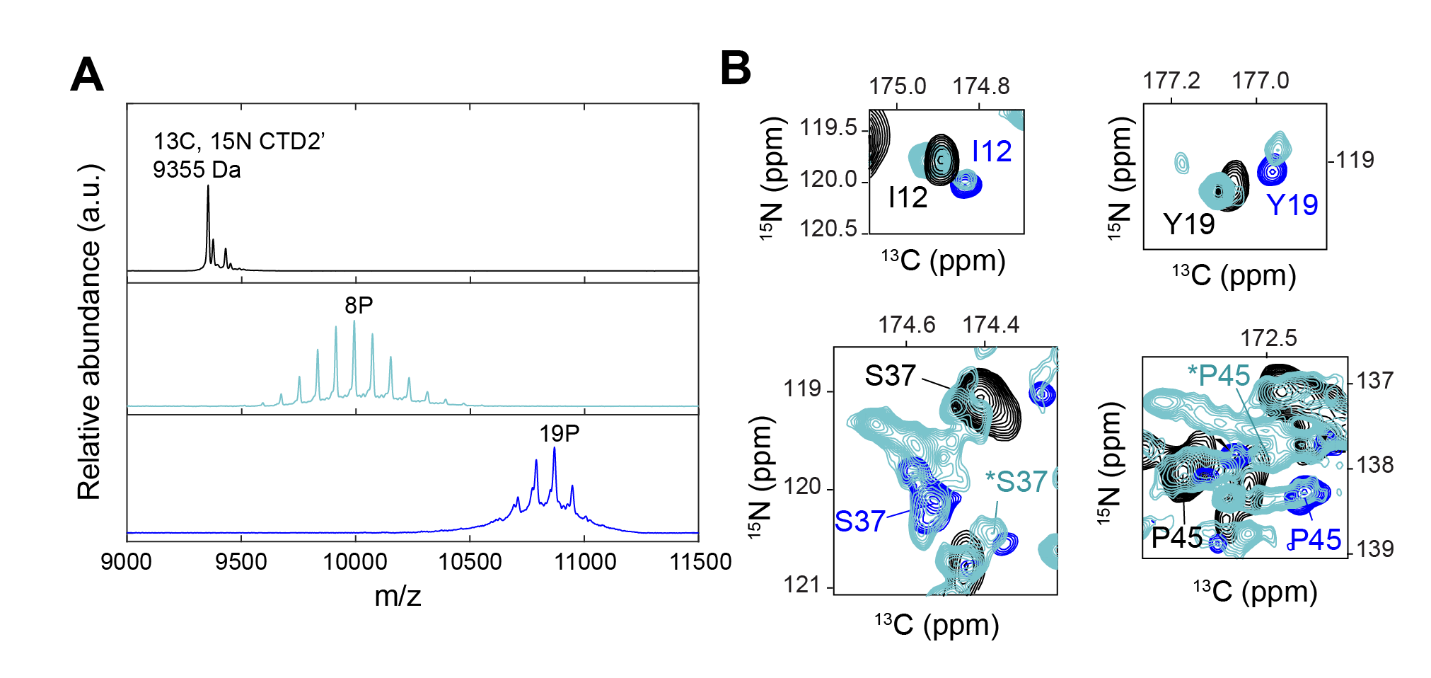

**Figure S2. NMR on Dyrk1a phosphorylated CTD2’. (A)** MALDI-TOF mass spectra of unphosphorylated, and two Dyrk1a phosphorylated ^13^C, ^15^N CTD2’ NMR samples. **(B)** Representative peaks from CON spectra showing Dyrk1a phosphorylated CTD2’ 8P (cyan) overlapping with unphosphorylated CTD2’ (black) and hyperphosphorylated CTD2’ 19P (blue), as well as unique peaks for Dyrk1a phosphorylated CTD2’ 8P denoted by asterisks (*).

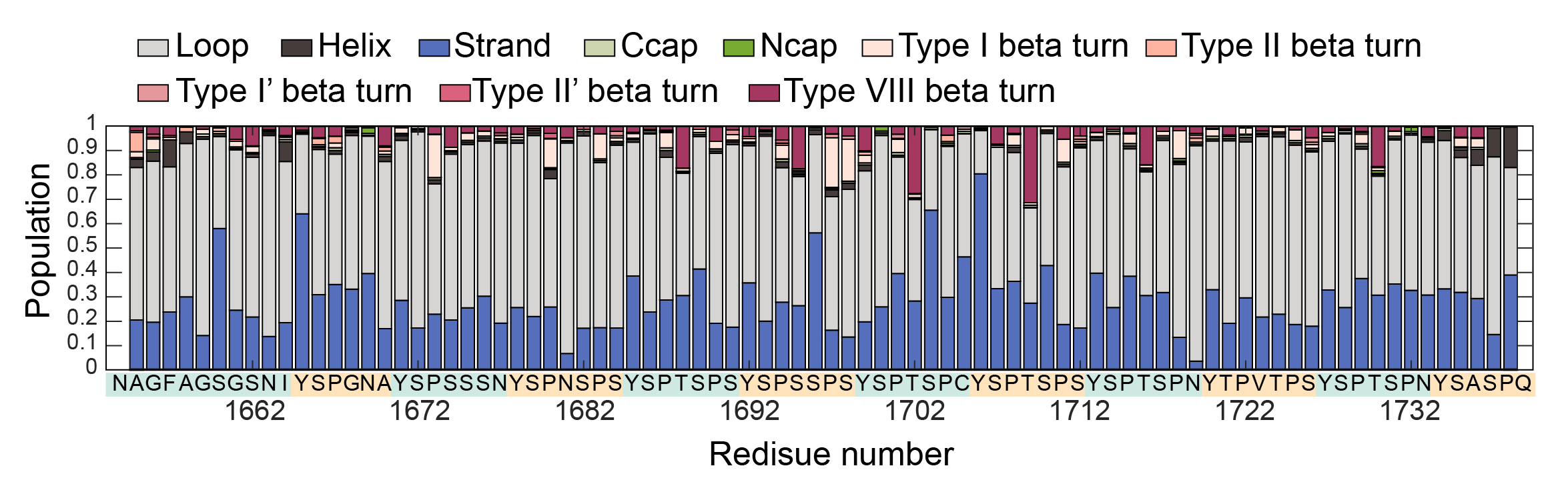

**Figure S3. Predicted structural motif populations in unphosphorylated CTD2’ using the MICS (Motif Identification from Chemical Shifts) program**. Populations of loops, helices, strands, N-terminal and C-terminal helix capping motifs (Ncap and Ccap), and five types of beta-turns (I, II, I’, II’, and VIII) were calculated based on backbone and sidechain chemical shifts.

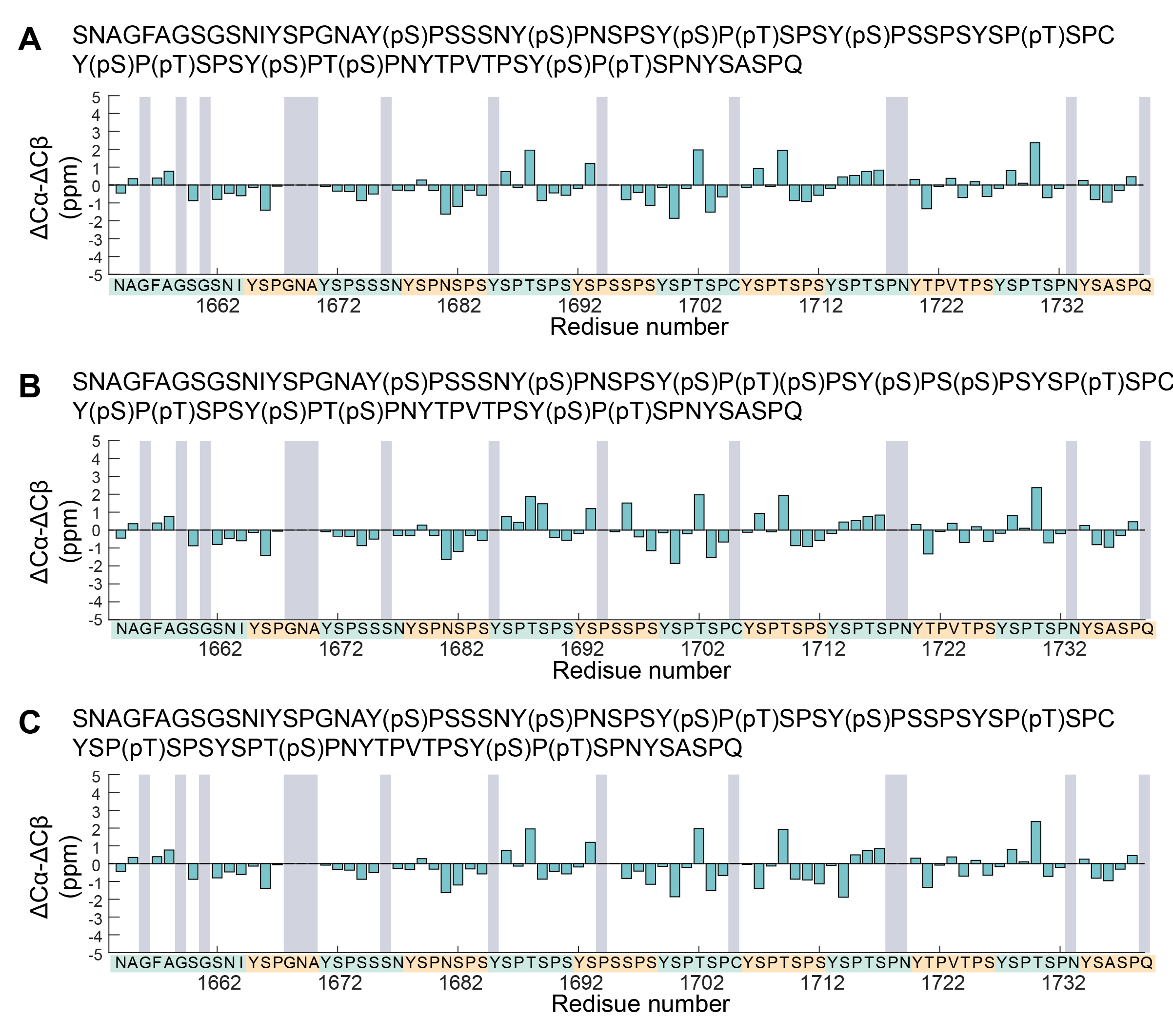

**Figure S4. Secondary structure propensity (SSP) for Dyrk1a phosphorylated CTD2’ (8P) (A–C)** SSP generated using three different random coil reference sets from different phosphorylation assignments. Residues without available Cα and/or Cβ chemical shift values are indicated by grey areas.

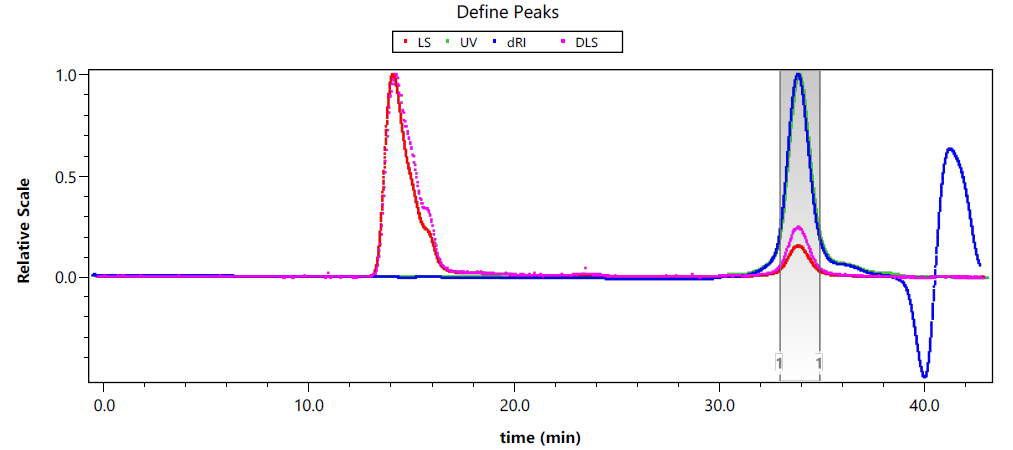

**Figure S5. SEC-MALS on CTD2’.** CTD2’ eluted at ~34 min as a monomer with a molecular weight of 8845 (±1.507%) Da, consistent with its theoretical molecular weight 8848 Da.

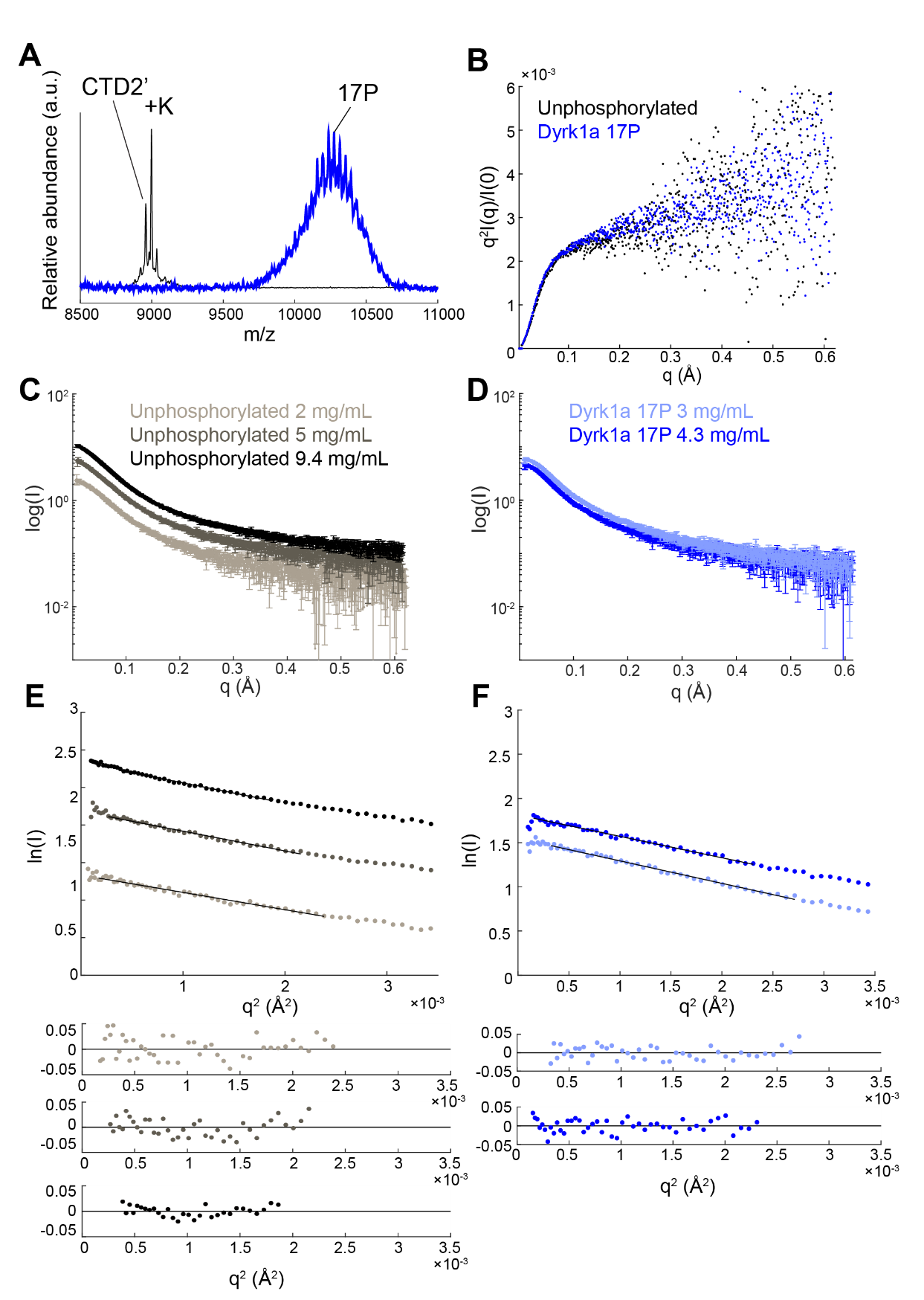

**Figure S6. SAXS on unphosphorylated and Dyrk1a phosphorylated CTD2’.** **(A)** MALDI-TOF mass spectra showing the SAXS sample of Dyrk1a phosphorylated CTD2’ contained an average of 17 phosphorylation marks. **(B)** Kratky plots for unphosphorylated and Dyrk1a phosphorylated CTD2’. **(C)** Raw scattering data for unphosphorylated CTD2’. **(D)** Raw scattering data for Dyrk1a phosphorylated CTD2’. **(E)** Guinier fitting and residuals for unphosphorylated CTD2’. **(F)** Guinier fitting and residuals for Dyrk1a phosphorylated CTD2’.

**Table S1. SAXS analyses on unphosphorylated and Dyr1ka phosphorylated CTD2’**

| **Unphosphorylated CTD2’** | | | | | | |
| --- | --- | --- | --- | --- | --- | --- |
| Concentration (mg/mL) | R_g_ (Å) Guinier | Range (points) | I(0) | R_g_ (Å) from P(r) | Range (points) | Dmax (Å) |
| 2 | 26.13± 0.31 | 13-57 | 2.29 ± 0.017 | 28.42 ± 2.34 | 13-368 | 115 |
| 5 | 27.99 ± 0.28 | 17-54 | 5.36 ± 0.035 | 33.55 ± 5.67 | 17-341 | 135 |
| 9.4 | 28.48 ± 0.22 | 21-50 | 10.28 ± 0.052 | 31.58 ± 10.56 | 21-335 | 120 |
| **Dyrk1a phosphorylated CTD2’** | | | | | | |
| Concentration (mg/mL) | R_g_ (Å) Guinier | Range (points) | I(0) | R_g_ (Å) from P(r) | Range (points) | Dmax (Å) |
| 3 | 28.24 ± 0.25 | 11-53 | 4.76 ± 0.027 | 31.49 ± 4.49 | 11-337 | 128 |
| 4.3 | 26.88 ± 0.22 | 12-56 | 6.12 ± 0.032 | 28.95 ± 6.26 | 12-355 | 110 |
